# Can machine learning aid in identifying disease genes? The case of autism spectrum disorder

**DOI:** 10.1101/2020.11.26.394676

**Authors:** Margot Gunning, Paul Pavlidis

**Affiliations:** Michael Smith Laboratories, University of British Columbia, Vancouver BC, V6T 1Z4, Canada; Department of Psychiatry, University of British Columbia, Vancouver BC, V6T 1Z4, Canada; Graduate Program in Bioinformatics, University of British Columbia, Vancouver BC, V6T 1Z4, Canada; Djavad Mowafaghian Centre for Brain Health, University of British Columbia, Vancouver BC, V6T 1Z4, Canada

**Keywords:** guilt by association, machine learning, disease gene prioritization, autism spectrum disorder

## Abstract

Discovering genes involved in complex human genetic disorders is a major challenge. Many have suggested that machine learning (ML) algorithms using gene networks can be used to supplement traditional genetic association-based approaches to predict or prioritize disease genes. However, questions have been raised about the utility of ML methods for this type of task due to biases within the data, and poor real-world performance. Using autism spectrum disorder (ASD) as a test case, we sought to investigate the question: Can machine learning aid in the discovery of disease genes? We collected thirteen published ASD gene prioritization studies and evaluated their performance using known and novel high-confidence ASD genes. We also investigated their biases towards generic gene annotations, like number of association publications. We found that ML methods which do not incorporate genetics information have limited utility for prioritization of ASD risk genes. These studies perform at a comparable level to generic measures of likelihood for the involvement of genes in any condition, and do not out-perform genetic association studies. Future efforts to discover disease genes should be focused on developing and validating statistical models for genetic association, specifically for association between rare variants and disease, rather than developing complex machine learning methods using complex heterogeneous biological data with unknown reliability.

## Introduction

Elucidating the genetic architecture of complex human disorders and diseases is currently a major challenge in medical research. Identifying genes involved in disease is often a time consuming and expensive process, so many researchers have been attracted to the idea of using predictions generated by machine learning (ML) algorithms (Krishnan et al., 2016; Lee et al., 2011; Moreau & Tranchevent, 2012; Y. Zhang et al., 2020). However, the effectiveness of ML approaches, in contrast to traditional genetic association, is unclear.

Algorithms used in gene function or disease prioritization tasks generally operate on a principle called guilt by association (GBA) (Gillis & Pavlidis, 2011b; Lanckriet et al., 2004), which postulates that genes with “associations” are more likely to be “guilty” of sharing functions. Associations can be sourced from multiple data types, such as gene expression, physical or genetic interactions, and protein sequence similarity. There are many ways these data types can be integrated into a machine learning method, and depending on the data types and algorithm, the associations among genes may be implicitly or explicitly represented as a network in which both direct and indirect associations can be used for inference.

Previous work from our group has shown that applications of ML to gene function prediction are highly influenced by biases in the underlying data (Gillis & Pavlidis, 2011, 2012; Pavlidis & Gillis, 2012). For example, protein interactions are often biased toward well studied genes, which often have high numbers of associated functional annotations (“multifunctional”). Furthermore, annotations and number of associations can be correlated, and this turns out to be a driver of GBA behavior: GBA tends to ascribe new functions to genes which are highly connected within the network rather than learning additional, novel information from the connection patterns (Gillis & Pavlidis, 2011; Pavlidis & Gillis, 2012). The implication of this “multifunctionality bias” is that GBA can seem to work in cross-validation settings, while providing predictions with little specific value. As an extreme illustration of this phenomenon, a million-edge network of gene associations can be reduced to 23 associations while not substantially impacting GBA performance (Gillis & Pavlidis, 2012). For these and other reasons, the real-life performance of GBA methods can be questioned. Focusing on the disease gene identification case, we are not aware of any instances where GBA has been responsible for a *bona fide* disease gene identification.

Recently there has been interest in a specific use case for GBA-based ML: predicting genes responsible for genetic risk of autism spectrum disorder (ASD). Multiple GBA-based ML studies have been produced with claims of providing greater insight into the genetic etiology of ASD (Brueggeman et al., 2020; Duda et al., 2018; Krishnan et al., 2016; Lin et al., 2018; Y. Zhang et al., 2020). ASD is a neurodevelopmental disorder with a genetically heterogenous etiology (de la Torre-Ubieta et al., 2016). Currently, much ASD research is aimed at identifying very rare, highly penetrant *de novo* variants in ASD probands because this class of variation has been found to impart a large proportion of risk (Iossifov et al., 2012; Neale et al., 2012; O’Roak et al., 2012; Sanders et al., 2012). While statistical methods for evaluating rare variants are still a topic of active research, genetic association underpins all ASD risk gene identification to date. In this context, ML methods have a challenge for acceptance by geneticists. Assessing the quality of ML-based ASD gene predictions is essential to provide realistic estimates of performance or complementarity to genetic approaches. However, we are unaware of attempts to compare them directly to genetic association studies, and assess their real-world applicability. In this paper we aimed to assess the reliability and usability of guilt by association machine learning approaches for ASD gene prioritization.

## Methods

### ASD gene sets

We compiled two ASD genes sets for algorithm evaluation (Table 1). We used the SFARIGene 2.0 (Abrahams et al., 2013) database as a source of well known, high-confidence ASD genes. SFARIGene collects information on ASD genetic risk factors and genes, and is manually curated by MindSpec, Incorporated. Genes are categorized by amount and quality of evidence for associated with ASD, and assigned a score ranging from 1 (high confidence) to 3 (suggestive evidence), or S (syndromic). We used the 144 genes from SFARI category 1 currently considered to be high-confidence ASD genes (Feb 2020) as our SFARI-HC gene set. Many of SFARI-HC genes were initially identified by the genetic association studies of De Rubies et al. (2014) and Sanders et al. (2015). Different subsets of these high-confidence genes had been used for training of the GBA ML algorithms discussed below. We complied a second high-confidence ASD risk gene set recently identified in three large-scale Transmission and *De Novo* Association Analysis (TADA) studies: Ruzzo et al. (2019) (iHart), Feliciano et al. (2019) (Spark), and Satterstrom et al. (2020). These three studies were built based on the background of the original TADA genetic association studies, De Rubeis et al. (2014) and Sanders et al. (2015). We refer to this set as “novel-HC” to reflect that most of the genes on this list were not used in the training of the GBA ML algorithms, largely because they were identified after the publication of the ML methods. We considered evaluation using the novel-HC genes a “testing scenario” because the ultimate use case of the machine learning algorithms is to highly prioritize and predict novel ASD genes.

**Table 1:**
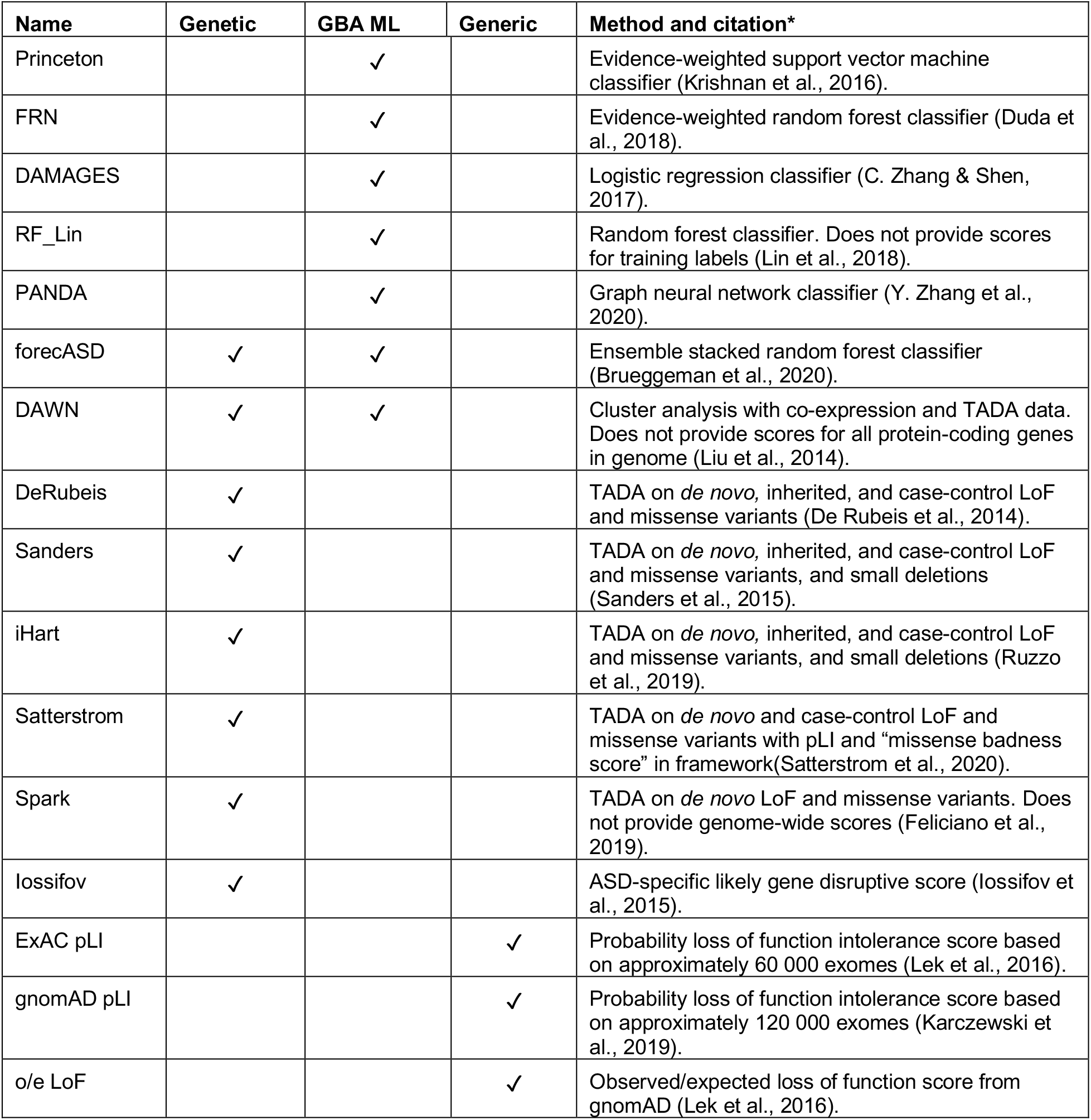
Summary of the ASD gene prioritization studies and generic methods for disease gene prioritization we used.

### ASD gene prioritization studies and generic measures of disease gene likelihood

We considered thirteen ASD gene prioritization studies (Table 1). Each study scored genes based on the authors’ assessment of their probability of contributing to ASD risk. All studies also provided lists of genes they considered to be high-confidence ASD risk gene candidates based on a thresholding of their rankings. We obtained these scores from the supplemental tables of the publications. We also evaluated three measures of constraint against loss-of-function (LoF) variation because they can be thought of as generic measures of disease gene likelihood (see below for descriptions).

We mapped gene symbols and Entrez gene identification numbers provided by each study to NCBI official gene symbols and Entrez gene identification numbers, and kept only protein-coding genes (Sayers et al., 2019). We used the mean score when a gene was listed more than once in a study. We ranked the scores from each study so that 1.0 was the highest possible score, indicating higher assessed likelihood of being involved in ASD, and 0.0 was the lowest possible score. The probability loss of function scores (pLI) from ExAC and gnomAD were already in the proper scale, with higher scores indicating genes likely to have high constraint against LoF variation. The scale of the observed/expected LoF score is opposite to the pLI scale and does not range from 0-1. We ranked genes based on o/e LoF score from lowest to highest. Lastly, for protein-coding genes not assessed in each GBA ML and GA study, we set the prediction or association score to be 0.0, or in the case of the o/e LoF score, the highest observed value of 2.0. Studies are organized into four categories based on the approach they used. Below we provide a brief description of each data source; see Supplemental Materials for more information.

### Genetic association studies

The studies described below are among the most important in terms of identifying what are generally considered high-confidence ASD genes (De Rubeis et al., 2014; Feliciano et al., 2019; He et al., 2013; Ruzzo et al., 2019; Sanders et al., 2015; Satterstrom et al., 2020). We included them in this study primarily to help establish a baseline to which the GBA ML approaches described in subsequent sections can be compared. Many of these studies are based on the TADA approach.

*DeRubeis* (De Rubeis et al., 2014) used whole-exome sequencing (WES) data from approximately 13 000 samples from trios and case-controls to identify *de novo* and inherited LoF variants, and *de novo* likely damaging missense variants (Mis3 by PolyPhen2). They used a TADA analysis to identify 33 ASD risk genes at FDR < 0.1. Samples from the Autism Sequencing Consortium (ASC), from Simons Simplex Consortium (SSC) (O’Roak et al. (2012), Sanders et al. (2012), and Iossifov et al. (2012)), and other cohorts were used. Association scores were provided for 18,735 genes.

*Sanders* (Sanders et al., 2015) used WES data from approximately 17,000 samples from trios and case-controls to identify *de novo* and inherited LoF variants, *de novo* likely damaging missense variants (Mis3 by PolyPhen2), and small *de novo* deletions. They employed a TADA analysis to identify 65 ASD risk genes at FDR < 0.1. They sequenced roughly 2 500 SSC families in addition to using SSC samples from Levy et al. (2011), Iossifov et al. (2014) and Dong et al. (2014), and ASC samples from De Rubeis et al. (2014), and samples from Pinto et al. (2014), among others. Association scores were provided for 18 665 genes.

*iHart* (Ruzzo et al., 2019) used whole-genome sequencing (WGS) data from 2 308 individuals from 493 multiplex Autism Genetic Resource Exchange (AGRE) families to identify *de novo* and inherited LoF variants and *de novo* likely damaging missense variants (Mis3 by PolyPhen2). They used their data and the Sanders data, and the Sanders TADA model to identify 69 ASD risk genes with FDR < 0.1, including 16 novel findings. Association scores were provided for 18,472 genes.

*Spark* (Feliciano et al., 2019) was the pilot study for the Simons Powering Autism Research for Knowledge (SPARK) project. They identified inherited and *de novo* likely damaging missense mutations (CADD >= 25) in 465 SPARK trios. They combined their *de novo* variants with *de novo* variants from 4 773 other simplex ASD trios from the ASC (De Rubeis et al. (2014)) and SSC (Iossifov et al. (2104); Krumm et al. (2015)), among other sources, for a TADA analysis. They identified 67 genes with FDR < 0.1, with 13 novel findings. They provided scores for the 2,249 genes found to have additional variation in SPARK families (Feliciano et al., 2019).

*Satterstrom* (Satterstrom et al., 2020) is the most recent and largest-scale genetic association study, with over 30 000 samples. They used samples from the SSC (Iossifov et al. (2012); Iossifov et al. (2014); O’Roak et al. (2012); Sanders et al. (2012)), the ASC (De Rubeis et al. (2014) and others), others from the AGRE and many other cohorts around the world. They used WES to identify *de novo* and case-control LoF, and *de novo* missense mutations (predicted by MPC, the “missense, PolyPhen-2, constraint score”), and employed TADA analysis to identify 102 ASD risk genes at FDR < 0.1. They considered 31 significant genes to be novel findings. Association scores were provided for 17,484 genes. Importantly, Satterstrom et al. modified the TADA method from the studies mentioned above by using the pLI score from ExAC and the MPC score to estimate the priors for the relative risk of LoF and missense variant classes.

*Iossifov* (Iossifov et al., 2015) computed a “Likely Gene-Disruptive” (LGD) score based on recurrence of LGD variants, the difference in frequency of LGD variants between ASD probands and unaffected siblings (ascertainment differential), and the load of LGD variation in ASD probands. They used data from WES of 2,471 families from the SSC (Iossifov et al. (2014)), and exome variants from approximately 6,000 controls from the Exome Variant Server (Iossifov et al., 2015). The theory behind the LGD score is similar to the TADA test and to generic measures of constraint against LoF and missense variation because they use recurrence of variants across multiple samples and models of expected LGD variation in a typical gene to increase power to find disease genes (He et al., 2013; Iossifov et al., 2015; Lek et al., 2016). They provided scores for 23,953 genes, and identified their top 239 genes as likely ASD risk gene candidates (Iossifov et al., 2015).

### GBA ML studies

Studies in this class do not use information from ASD genetic association studies, but they use machine learning algorithms to distinguish ASD from non-ASD risk genes using other types of non-genetics data.

*Princeton* (Krishnan et al., 2016) is an evidence weighted support vector machine (SVM) built on a functional interaction network made from human gene expression, protein-protein interaction, regulatory, and genetic and chemical perturbation data. For training they used 594 ASD genes, and 1,189 manually curated non-mental health associated genes as positives and negatives, respectively. The positive ASD genes were given one of three weights (1.0, 0.5, 0.25) based on strength of evidence of association with ASD. Krishnan et al. provided likelihood rankings for 25 825 genes, and identified their top decile as likely ASD risk gene candidates.

*FRN* (Duda et al., 2018) is a random forest classifier built on an evidence-weighted functional interaction network of human, mouse and rat brain gene expression, protein-protein interaction, protein docking and phenotype annotation data. They used 143 high-confidence ASD genes from SFARI and the Sanders publication above as positive training genes, and 1,176 of the of the 1,189 Princeton non-mental health associated genes as negative training genes. They provided likelihood rankings for 21,114 genes, and identified their top decile as likely ASD risk gene candidates.

*DAMAGES* (C. Zhang & Shen, 2017) used a combination of regularized and logistic regression using cell-type specific gene expression data and measures of constraint against LoF and missense variation from ExAC. First, they created a DAMAGES (D) score using principal component analysis (PCA) and regression analysis on gene-expression profiles of 24 mouse central nervous system cell types in 6 regions. They created profiles of 145 genes found to have *de novo* LoF variants in ASD probands and unaffected siblings from multiple cohorts as training samples. Next, using logistic regression, they combined the D score with ExAC measures to create an ensemble (E) score. They used 36 genes with 2 or more *de novo* LoF variants in ASD probands, and 156 genes with 1 or more *de novo* LoF variants in sibling controls from multiple cohorts as positive and negative training genes, respectively. They provided likelihood rankings for 15,881 human genes, and identified their top 117 genes as likely ASD risk gene candidates.

*RF_Lin* (Lin et al., 2018) is a random forest classifier. They built an evidence-weighted network of BrainSpan co-expression and protein-protein interaction data, and extracted network features such as hubness and centrality (Sunkin et al., 2013). The features of their classifier included their network association matrix, selected network features, and gene-level constraint measures from ExAC. They used the positive and negative training labels employed by FRN described above. They provided likelihood rankings for 17 099 genes, and identified their top decile as likely ASD risk gene candidates. They did not provide scores for their training genes.

*PANDA* (Y. Zhang et al., 2020) used a network-based deep-learning approach to prioritize autism genes. They built a human molecular interaction network from protein-protein interaction data from multiple sources, and used a training set of 760 ASD genes from SFARI Gene 2.0 and OMIM weighted by confidence of association with ASD (1.0, 0.75, 0.5). They provided likelihood rankings for 23,472 genes, and defined an “autism subnetwork” made up of 2,346 genes (approximately top decile).

### Genetics-GBA Hybrid ML studies

The studies in this section used a combination of ASD-specific genetic association information (e.g. from the studies listed above) along with other features to build their models. Information from the two classes of features are integrated prior to training a machine learning algorithm to distinguish ASD from non-ASD risk genes, using high-confidence ASD genes from genetic association studies as their positive training set.

*DAWN* (Liu et al., 2014) built a co-expression network from BrainSpan data of the prefrontal and motor-somatosensory neocortex at 10-24 weeks post-conception, and overlaid association statistics from a TADA analysis (Sunkin et al., 2013). Using unsupervised model-based clustering (Weighted Gene Co-expression Network Analysis) and a hidden Markov random field, they modeled the correlation of genetic association scores across the co-expression network to identify highly correlated nodes, or “network ASD genes.” Following a false discovery rate estimation procedure, they identified 127 likely ASD risk gene candidates from 10,233 genes.

*forecASD* (Brueggeman et al., 2020) is a stacked random forest ensemble classifier using BrainSpan(Sunkin et al., 2013) gene expression data, STRING(Szklarczyk et al., 2019) protein-protein interaction data, and genome-wide results from Princeton, DAWN, DAMAGES, Sanders and DeRubeis studies described above. They used 76 SFARI high-confidence genes and 1,000 randomly selected non-SFARI genes as positive and negative training examples, respectively. They provided likelihood rankings for 17 957 genes, and identified their top decile of genes as likely ASD risk gene candidates.

### Generic measures of disease gene likelihood

The scores in this section were developed without any disease specificity, and measure the depletion of LoF variation within a gene. Therefore, these scores act as generic proxies for the likelihood of a gene to be involved in *any* genetic disease. We downloaded these scores from the gnomADv.2.1.1 database on 2019-07-18 (Karczewski et al., 2019).

*ExAC_pLI* measures the probability of a gene to be extremely intolerant of LoF variation. It’s scale is ranges from 0-1, with genes over 0.9 representing those extremely intolerant to LoF variation and under higher constraint. It was developed based on data from approximately 60,000 exomes. *GnomAD_pLI* is similar but computed from an expanded data set of roughly 120,000 exomes.

*oe_LoF* measures the deviation of the number of observed LoF variants within a gene to the expected number. This score differs from the above two because its scale is reversed, with scores below 0.35 indicating extreme depletion of LoF variation and higher constraint (Karczewski et al., 2019). This measure was recommended for identifying genes likely to be depleted of LoF variation because it is more interpretable than the pLI (i.e. a score of 0.4 indicates that 40% of the expected LoF variants within a gene have been observed), and better captures intermediate levels of happloinsufficiency. In addition, ynlike the pLI, the o/e LoF score reports a confidence interval; the upper 95% confidence bound is recommended as the criterion to be compared to 0.35 (Karczewski et al., 2019).

### Evaluation

We plotted receiver operating characteristic (ROC) curves and precision-recall curves to assess recovery of the novel high-confidence (novel-HC) and SFARI high-confidence (SFARI-HC) ASD gene sets. When evaluating the ability of the scores to rank the novel-HCASD gene set, we removed the SFARI high-confidence ASD genes and other ASD genes used in the training of the ML algorithms from their gene rankings. This was done to ensure that the algorithms were not penalized for performance on ASD genes. The top ranks provided by the studies are their predictions as to the most likely ASD risk candidates. Therefore, the PR curves are the preferred evaluation metric because they are more sensitive to classification errors in the top ranks.

We calculated area under the ROC curve (AUROC) using the “auc” function with the “trapezoid” method from the DescTools R package (Andri Signorell et mult. al., 2019) to account for ties in the rankings. We calculated precision at 20% recall (P20R) of total genes in the ASD gene sets, and precision at 43% recall (P43R) of total genes in the ASD gene sets. Precision at 20% recall was selected as a ‘midrange’ for display purposes, and has previously been used as a reported point statistic in function prediction algorithm assessments (Peñ a-Castillo et al., 2008). The exception is for the pLL scores; as more than 20% of the high-confidence ASD genes have the maximum pLI score of 1.0, we report precision at 43% recall to have a consistent comparison for precision-recall across all studies.

We used 2,500 bootstrapped samples (gene-level) to calculate 95% confidence intervals for AUROC and precision-recall statistics. The bootstrapped samples were stratified, and done with replacement. This means that we sampled from the ASD gene sets and the rest of the scored protein coding genes separately in each of the 2,500 iterations to ensure balanced coverage, and that the same gene could be sampled more than once in each iteration. Therefore, in each bootstrapped sample, we kept only unique genes for evaluation. Studies whose performance measures confidence intervals did not overlap were considered significantly different from each other.

We measured the correlation between the ASD gene likelihood rankings provided by each study, and other metrics of interest using the Spearman correlation coefficient. The other metrics of interest included a multifunctionality rank, node degree of a BioGrid protein-protein interaction network, number of publications, and the SFARI numeric gene score. If a gene did not have a score for other metrics of interest, it was given a value of 0.0 for consistency with ASD gene prioritization studies.

Each method provided a cut off for their set of likely ASD genes, and we calculated the overlap in their top gene sets as their shared number of genes.

### forecASD analysis

We obtained code for the forecASD classifier from https://github.com/LeoBman/forecASD, and re-ran it locally (Brueggeman et al., 2020). A minor difference from the preprint is the GitHub code uses a different version of the randomForest R package (version 4.6-14 vs 4.6-12 in the preprint) (Brueggeman et al., 2020; Liaw & Wiener, 2002). We refit the final ensemble model (03_ensemble_model.R) with different sets of the input features used in final ensemble model: the noClass (noC) model removed features from other classifiers listed above; the noClassPPI (noCP) model eliminated the other classifiers, and the STRING score; noClassPPIBS (noCPB) model eliminated the other classifiers, the STRING score, and the BrainSpan score; the PPIOnly (PPI) model only used the STRING score; and the BrainSpanOnly (BS) model only used the BrainSpan score. Feature importance was measured by mean decrease in accuracy and mean decrease in Gini node impurity. Mean decrease in accuracy is measured by randomly permuting each feature, and measuring the out-of-bag (cross-validation) accuracy of the resulting trees. Mean decrease in Gini measures how well the features can split the data from mixed labelled nodes into pure single class nodes. Brueggeman et al. did not provide code for their feature importance plots; we used “varImpPlot” from the randomForest package (Brueggeman et al., 2020; Liaw & Wiener, 2002). When rerunning their provided code, we found that two columns in their metadata had been mislabelled, D (DAMAGES) and D_ens (DAMAGES ensemble), necessitating re-labelling for plotting of feature importance. As for the other methods, we evaluated each model using the two ASD gene sets and with the same metrics described above.

## Results

### Method outline

We considered thirteen ASD gene prioritization studies, and three measures of generic disease gene likelihood for evaluation. Each study provided scores for genes based on the author’s assessment of their probability of contributing to ASD risk. We evaluated their ability to prioritize novel high-confidence and known high-confidence ASD genes using ROC and Precision-Recall curves, and 95% confidence intervals of AUROC and precision at 20% recall. Additionally, we looked at how the scores correlated with one another, and with other generic network features such as number of physical interaction partners to assess potential biases.

### Systems–based GBA ML methods do not prioritize novel high-confidence ASD genes well compared to other disease gene prioritization methods

The first test we performed was investigating how well the GBA ML studies prioritized novel high-confidence ASD genes which were not used to build their predictions, and comparing their performance to genetics-based and generic approaches for disease gene prioritization. Because genetic association remains the gold standard method for identifying genetic risk factors, our operating assumption was that in order to be considered a successful method, an ASD-specific GBA ML study should have comparable performance to the genetic association studies alone, and should outperform generic measures of disease gene likelihood. Lastly, the more recent genetic association studies (iHart and Satterstrom) were built up from the DeRubeis and Sanders studies in that they are using overlapping samples, and similar model parameters and variant classes in their TADA analyses. Therefore, we expected that the DeRubeis and Sanders studies would rank the novel-HC ASD genes at lower or borderline significant levels, and that the iHart and Satterstrom studies would show higher rankings of each other’s hits.

We found that GBA ML studies had comparable performance to the generic measures of disease gene likelihood, as is shown by their overlapping 95% confidence intervals for precision at 20% recall (i.e. P20R_FRN_ = 0.69-4.61%; P20R_ExAC_pLI_ = 0.69-1.92%) (Figure 1D, 1H, Table 2). While the studies had high AUROC statistics with overlapping 95% confidence intervals, these metrics are somewhat misleading because they are not sensitive to false positive predictions in top rankings, which are most relevant for prioritization studies (Figure 1B, 1F, Table 2). This finding suggests limited utility of GBA ML studies for ASD gene prioritization: use of a simple non-ASD specific measure constraint against LoF variation has comparable performance to complex ML approaches.

**Table 2:** Summary statistics on novel-HC genes. AUROC: Area under the receiver operator characteristic curve; P20R: precision at 20% recall; P43R: precision at 43% recall. Values in parentheses are the upper and lower 95% confidence interval bounds. A ‘*’ for P43R indicates a tie at recall of 20/43% of gene set.

| Paper | AUROC | P20R (%) | P43R (%) |
| --- | --- | --- | --- |
| Princeton | 0.67 (0.60, 0.74) | 0.88 (0.56, 1.00) | 0.85 (0.47, 1.16) |
| FRN | 0.73 (0.67, 0.79) | 1.28 (0.69, 4.61) | 0.87 (0.55, 1.33) |
| DAMAGES | 0.80 (0.75, 0.85) | 1.25 (0.87, 1.79) | 1.40 (0.76, 1.90) |
| RF_Lin | 0.77 (0.71, 0.83) | 2.12 (0.96, 3.94) | 1.49 (0.84, 1.87) |
| PANDA | 0.72 (0.66, 0.79) | 1.08 (0.44, 3.69) | 0.68 (0.43, 0.96) |
| forecASD | 0.89 (0.84, 0.94) | 7.13 (4.63, 11.21) | 5.56 (3.24, 9.38) |
| DAWN | 0.58 (0.53, 0.63) | 0.69 (0.37, 1.37) | 0.53 (0.35, 0.72)* |
| ExAC_pLI | 0.77 (0.72, 0.83) | 1.14 (0.69, 1.92) | 1.11 (0.78, 1.46) |
| gnomAD_pLI | 0.75 (0.7, 0.81) | 1.05 (0.70, 2.08) | 1.15 (0.72, 1.39) |
| oe_LoF | 0.79 (0.74, 0.84) | 0.96 (0.64, 1.45) | 1.10 (0.81, 1.37) |
| DeRubeis | 0.72 (0.65, 0.79) | 4.73 (2.08, 5.60) | 2.37 (1.44, 3.77) |
| Sanders | 0.76 (0.69, 0.84) | 6.20 (3.49, 10.54) | 2.80 (1.68, 6.33) |
| iHart | 0.78 (0.71, 0.85) | 42.35 (5.00, 58.33) | 4.07 (1.69, 7.61) |
| Satterstrom | 0.85 (0.79, 0.91) | 50.00 (37.57, 75.79) | 54.13 (24.41, 69.91) |
| lossifov | 0.76 (0.70, 0.82) | 1.82 (1.24, 3.65) | 1.83 (1.10, 2.68) |

**Figure 1:**
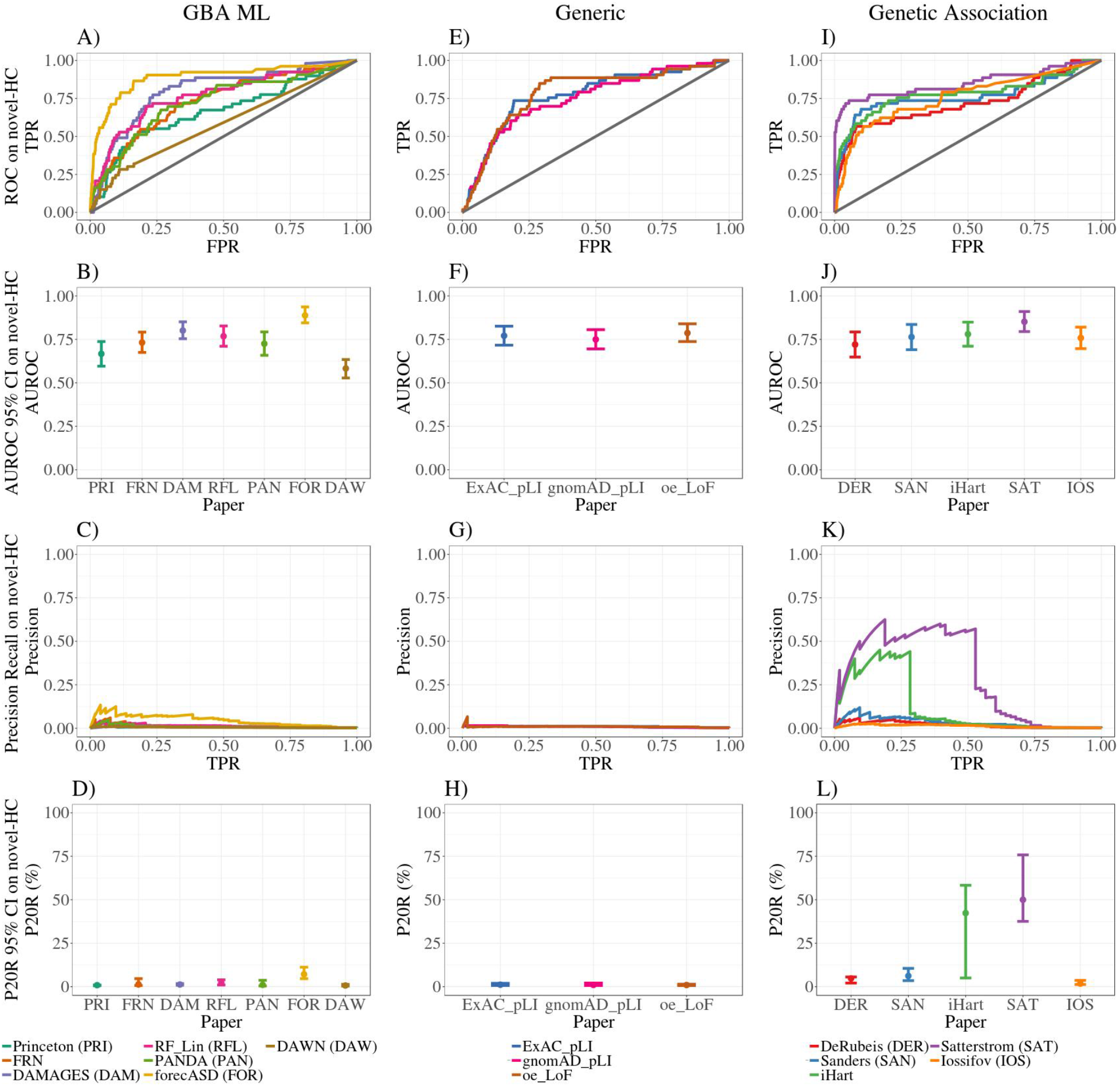
ROC, Precision-Recall and summary statistics on novel-HC genes. Novel-HC genes were discovered by new TADA studies (iHart, Spark and Satterstrom), and most were not used in training of GBA ML studies. GBA ML studies have comparable performance to generic measures of disease gene likelihood (LoF constraint measures), with high AUROC (A, B, E, F), but low precision at 20% recall (C, D, G, H). GBA ML methods incorporating genetics information, particularly forecASD, have significantly better performance. Note that DAWN does not provide likelihood estimates for all protein-coding genes in the genome. Genetic association studies also show high AUROC (I, J). Previous TADA studies (DER, SAN) show moderate performance while the newer TADA studies are not performing at 100% precision (L). 95% confidence intervals were created from 2 500 stratified bootstrap samples (B, D, F, H, J, L). TPR, True Positive Rate; FPR, False Positive Rate; AUROC, Area Under the Receiver Operator Curve; P20R, Precision at 20% Recall.

The best performing GBA ML method was the hybrid genetics-GBA method forecASD (P20R_forecASD_=4.63-11.21%), which had similar levels of performance to the genetic association studies developed before the iHart, Spark and Satterstrom studies (i.e. P20R_Sanders_=3.49-10.54%) (Figure 1D, 1L, Table 2). The other hybrid method, DAWN, has similar performance to other GBA ML studies, but this may be in part because they only provide predictions scores for roughly 10 000 genes in the genome (Figure 1A-D, Table 2).

It is important to note that the Satterstrom and iHart studies do not have particularly high performance by these benchmarks, despite being, in effect, a comparison of genetics findings to updated genetics findings (Figure 1L, Table 2). In other words, the two recent TADA-based studies do not agree on what genes are significantly associated with ASD. Additionally, the previous TADA studies have some performance (i.e. P20R_Sanders_=3.49-10.54%), which would suggest that they were able to identify some of the novel genes at marginal levels of significance, and with the accumulation of more data, these genes became significant in the newer studies (Figure 1L, Table 2).

### GBA ML methods do not predict high-confidence ASD genes

We analyzed how well the GBA ML studies recovered SFARI high-confidence genes, many of which were used in the training of the ML algorithms, and compared the results to other methods for disease gene prioritization. The genes in the SFARI-HC set were discovered by different genetic association studies, many of which were first identified by the DeRubeis and Sanders studies. Given the relationship between all the TADA studies, we expected the original and newer genetic association studies to highly prioritize SFARI high-confidence genes. The systems-based GBA ML studies used different subsets of SFARI high-confidence genes, and other ASD associated genes, during training. Therefore, we would expect that these studies should also highly prioritize SFARI-HC genes. We note that this is not a pure test of training performance because not all SFARI-HC genes were used during the training step. However, because the methods were developed at different times using different training gene sets, we opted for a consistent evaluation gene set across methods.

Our findings from this set of analyses parallel what we found for the novel-HC gene set. Mainly, we found that the GBA ML studies have comparable performance to the generic measures of disease gene likelihood with overlapping 95% confidence intervals for precision at 20% recall (i.e. P20R_FRN_ = 5.33-12.06%; P20R_ExAC_pLI_ = 9.68-14.08%) (Figure 2D, 2H, Table 3). RF_Lin did not provide predictions for their training genes, which partially explains its poorer performance relative to other studies (AUROC_RF_Lin_=0.44-0.55; P20R_RF_Lin_=3.49-7.08%) (Figure 2B, 2D, Table 3). Again we found that the genetics-GBA method forecASD had the best performance of the GBA ML studies with similar performance to genetic association studies (i.e. P20R_forecASD_ = 38.23-77.32%; P20R_Sanders_=53.74-95.00%; P20R_Satterstrom_=65.48-100.00%) (Figure 2D, 2L, Table 4). These results show that systems-based GBA ML studies are providing little ASD-specific information above that provided by the generic measures of constraint against LoF variation. Once again, these findings highlight the limited utility of the systems-based GBA ML studies for prioritizing ASD risk genes.

**Table 3:** Summary statistics on SFARI-HC genes. Headings are per table 2, and * means a tie at recall of 20/43% of gene set.

| Study | AUROC | P20R (%) | P43R (%) |
| --- | --- | --- | --- |
| Princeton | 0.81 (0.78, 0.84) | 6.14 (4.79, 7.58) | 4.36 (3.49, 6.02) |
| FRN | 0.88 (0.86, 0.90) | 8.00 (5.33, 12.06)* | 5.79 (4.53, 6.37) |
| DAMAGES | 0.87 (0.85, 0.90) | 7.36 (5.70, 8.90) | 7.13 (5.78, 8.78) |
| RF_Lin | 0.50 (0.44, 0.55) | 5.59 (3.49, 7.08) | 2.58 (0.53, 3.82) |
| PANDA | 0.87 (0.84, 0.89) | 7.67 (5.76, 11.99) | 5.78 (4.79, 6.80) |
| forecASD | 0.94 (0.92, 0.95) | 53.56 (38.23, 77.32) | 31.80 (23.85, 40.22) |
| DAWN | 0.62 (0.59, 0.65) | 3.00 (2.07, 7.62) | 1.70 (1.39, 2.02)* |
| ExAC_pLI | 0.88 (0.85, 0.91) | 12.06 (9.68, 14.08)* | 9.57 (8.31, 12.07)* |
| gnomAD_pLI | 0.88 (0.85, 0.91) | 11.42 (9.59, 12.96)* | 11.42 (9.29, 12.96)* |
| oe_LoF | 0.91 (0.89, 0.93) | 23.02 (13.80, 32.05) | 16.30 (12.29, 18.80)* |
| DeRubeis | 0.68 (0.63, 0.72) | 29.00 (11.89, 77.38) | 4.42 (1.68, 6.34) |
| Sanders | 0.73 (0.68, 0.77) | 85.29 (53.74, 95.00) | 12.45 (6.68, 20.99) |
| iHart | 0.73 (0.68, 0.77) | 74.36 (41.06, 97.22) | 14.28 (5.68, 28.05) |
| Satterstrom | 0.75 (0.70, 0.80) | 85.29 (65.48, 100.00) | 31.51 (16.37, 55.71) |
| lossifov | 0.88 (0.86, 0.91) | 56.85 (43.99, 68.00)* | 31.00 (18.04, 44.85) |

**Table 4:**
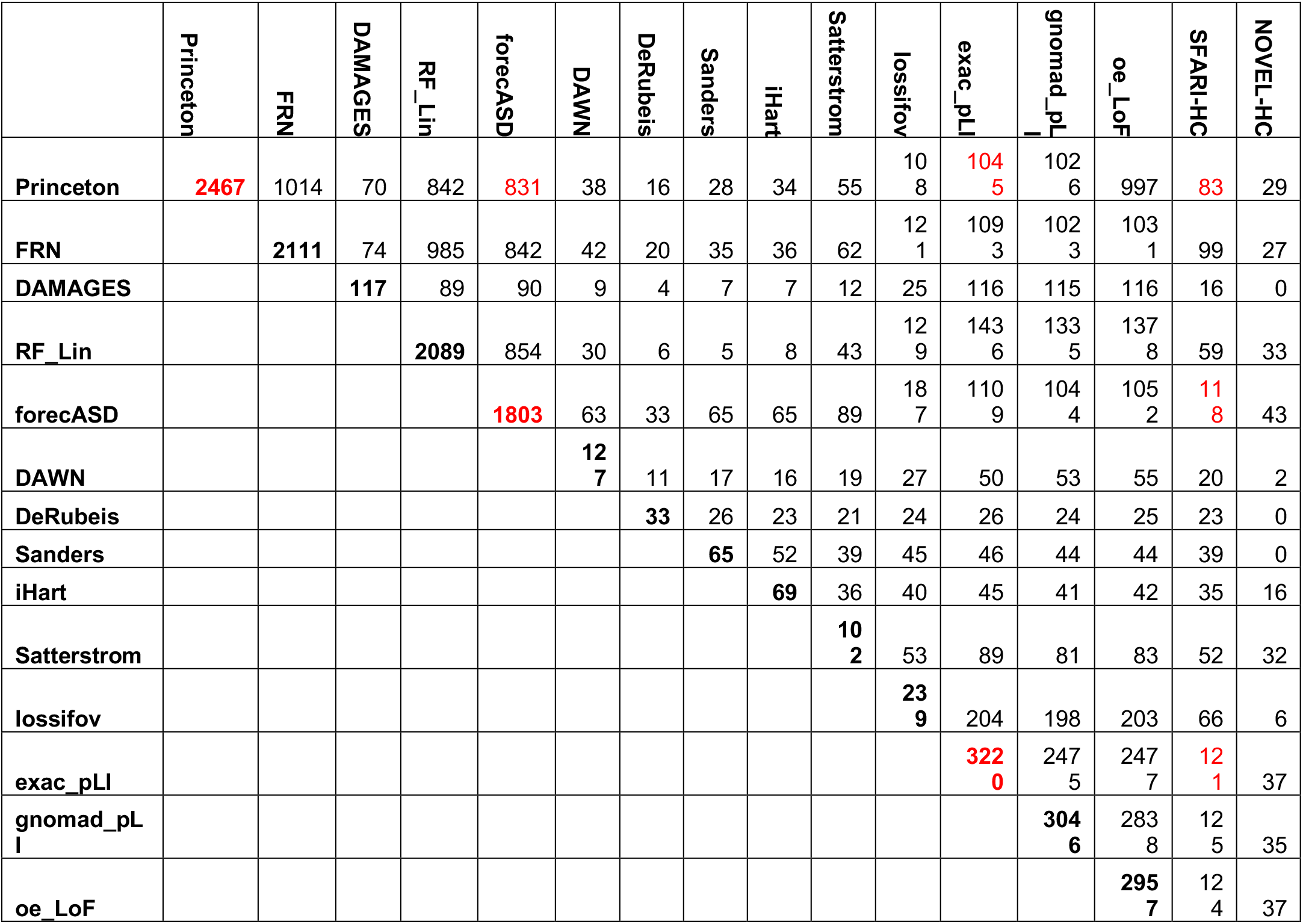
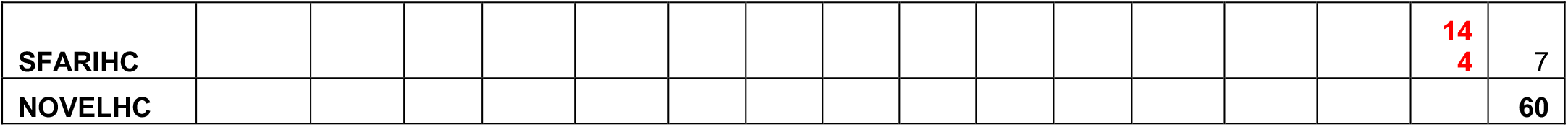
Overlap of top ranked ASD genes from each study. Numbers on the diagonal represent the number of ASD genes predicted. Values highlighted in red are discussed in the main text.

**Figure 2:**
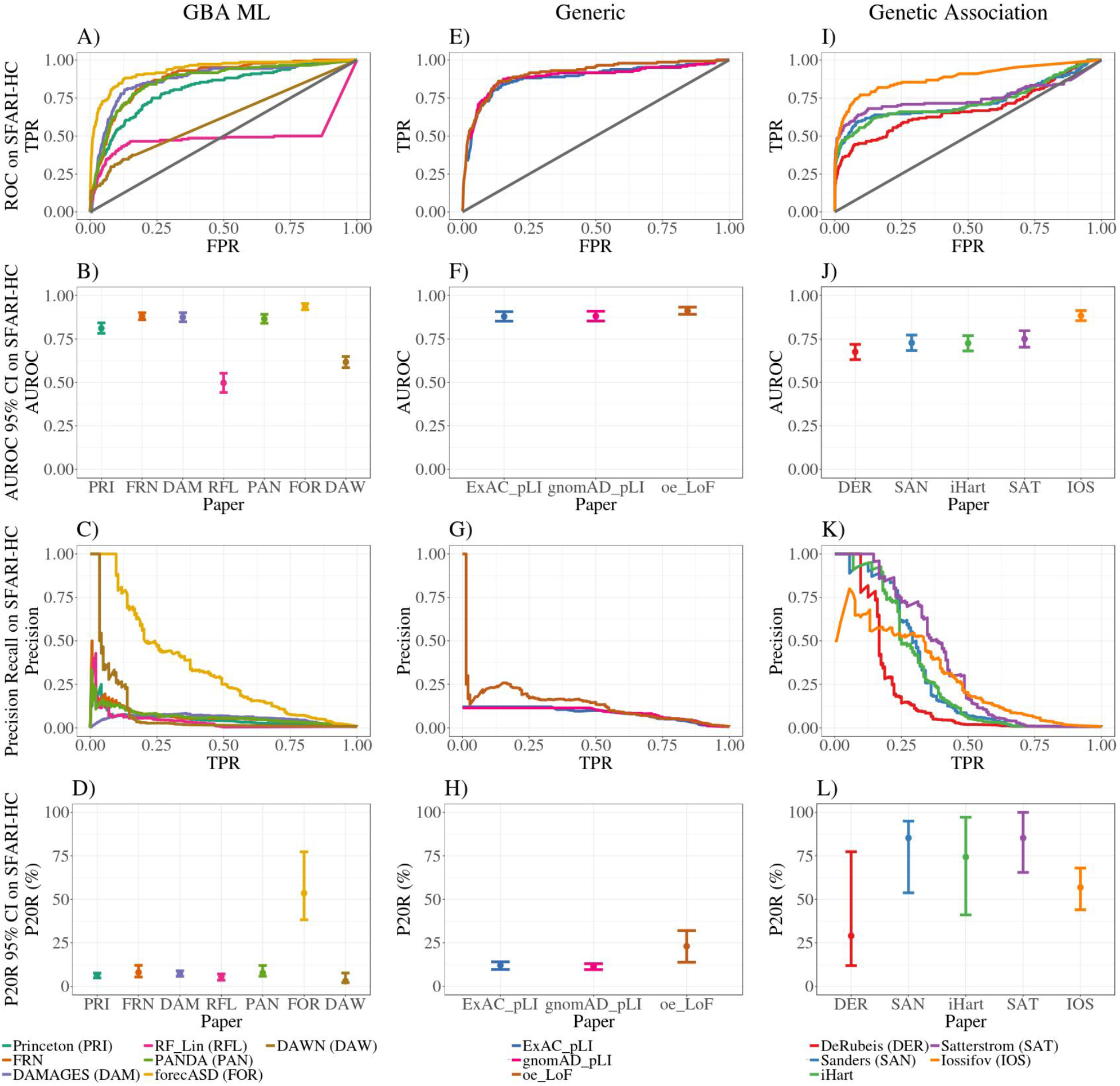
ROC, Precision-recall and summary statistics for SFARI-HC genes. Many SFARI-HC genes were initially discovered by early TADA studies (DER, SAN), and used in training of GBA ML studies. Thus, this evaluation acts as a control experiment. Systems-based GBA ML studies have comparable performance to generic measures of disease gene likelihood (LoF constraint measures), with high AUROC (A, B, E, F), but low precision at 20% recall (C, D, G, H) on SFARI-HC genes. The GBA ML method with genetics information, forecASD, had significantly better performance compared to other GBA ML methods. Note that DAWN does not provide estimates for all protein-coding genes in the genome, and RF_Lin does not provide estimates for their training genes, which partially explain their poorer performance. Genetic association studies show high AUROC (I, J) and high precision at 20% recall (K, L). 95% confidence intervals were created from 2500 stratified bootstrap samples (B, D, F, H, J, L). TPR, True Positive Rate; FPR, False Positive Rate; AUROC, Area Under the Receiver Operator Curve; P20R, Precision at 20% Recall.

### Low agreement between ML and genetic association

As previously discussed, GBA postulates that genes with shared associations are more likely to have shared functions or be involved in the same diseases. However, predictions can be driven by underlying multifunctionality bias whereby new functions are ascribed to genes that are well characterized because they are highly studied, and have a high number of association annotations (Gillis & Pavlidis, 2011; Pavlidis & Gillis, 2012). In other words, we hypothesized that GBA methods using heterogeneous biological networks biased towards well-studied genes would tend to rank generically “disease-related” genes highly simply because they are well studied. Furthermore, because this ranking is not ASD-specific, it cannot readily identify novel and specific relationships. On the other hand, methods which do not recapitulate these generic rankings may perform badly because the main source of apparent performance of GBA methods is their ability to prioritize well studied genes (“multifunctionality bias” as per Gillis and Pavlidis).

We compared the genetic association and GBA ML scores to generic network features and generic gene annotations (Figure 3). Our results show that some of the GBA ML studies are indeed biased. For example, the genetics-GBA study, forecASD, has moderate correlation with physical node degree (R_Spearman_ =0.34) and number of publications (R_Spearman_=0.34), as do DAMAGES, RF_Lin and PANDA (Figure 3). In the work of Gillis and Pavlidis, correlations of this magnitude were sufficient to explain a large fraction of predictive performance. In contrast, Princeton and FRN did not appear to show bias (i.e. R_S:FRN,pnd_=0.16, R_S:FRN,numPubs_=-0.03). Furthermore, as expected the TADA analyses show little to no agreement with these generic features (i.e. R_S:iHart,pnd_=0.05, R_S:iHart,numPubs_=0.05). These findings offer some explanation for the poor performance of the systems-based GBA ML studies when tested on novel genes. Methods which are not biased towards well studied genes, such as Princeton and FRN, may be performing poorly because there is no bias to drive apparent performance (Gillis & Pavlidis, 2011; Pavlidis & Gillis, 2012). On the other hand, studies which are biased towards well studied genes, such as RF_Lin and PANDA, may be performing poorly because GBA is assigning new functions to highly connected genes in the network, and not learning ASD-specific information (Gillis & Pavlidis, 2011; Pavlidis & Gillis, 2012). However, further work is required to delineate how multifunctionality is affecting each study. Lastly, high agreement between generic measures of constraint and systems-based GBA ML studies further suggests that their predictions are generic, and not specific to ASD (i.e. R_S:forecASD,ExAC_pLI_=0.37) (Figure 3).

**Figure 3:**
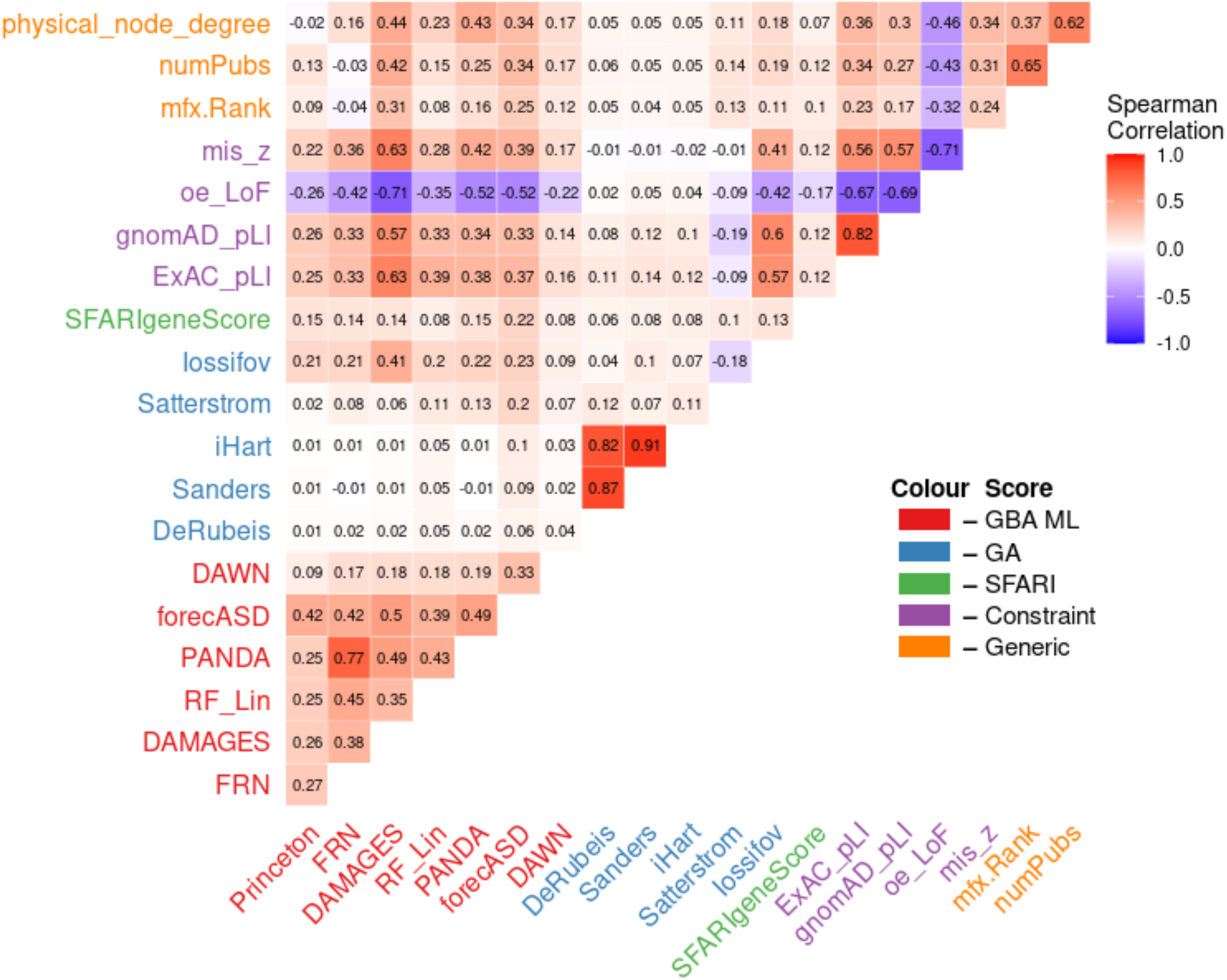
Correlations among gene rankings. Values are Spearman correlations. Notable patterns include low correlations between genetic association methods, ML methods and other network features such as node degree and publication number; increased correlation between select ML methods and other network features; low correlation between Satterstrom score and pLI despite its incorporation in the statistical framework; low correlation between SFARI gene score and generic gene annotations.

The lack of agreement between Satterstrom and the other TADA analyses further highlights the non-equivalence between the genetic association studies and the need for TADA model validation (i.e. R_S:Satterstrom, DeRubeis_=0.12) (Figure 3). Satterstrom directly incorporates ExAC pLI into its TADA model, however, it displays little correlation with pLI (R_S:ExAC_pLI_=-0.09), and low to moderate agreement with other generic network features (i.e. R_S:Satterstrom,numPubs_=0.14). While it is possible that using pLI incorporated some generic disease gene bias into Satterstrom, the direct effects of pLI on the score are likely complex and non-linear due to TADA’s approach of collapsing multiple pieces of information to derive the per-gene association scores (He et al., 2013; Satterstrom et al., 2020). Therefore, Spearman correlation may not adequately capture the relationship.

Iossifov is the genetic association study with the highest agreement with generic gene annotations. Notably, it has high correlation with pLI (R_S:ExAC_pLI_=0.60). Iossifov is the most similar to pLI in its construction: both scores attempt to quantity the deviation of the observed number of LoF variants from an expectation of LoF variation derived from complex models incorporating rates synonymous variation, among many other factors (Iossifov et al., 2015; Karczewski et al., 2019; Lek et al., 2016). The Iossifov score is ASD-specific because they incorporate an estimate of the number of causal ASD genes, and the observed load of LoF variation in ASD probands, whereas the LoF constraint scores were developed without any disease specificity (Iossifov et al., 2015).

Lastly, agreement between the SFARI gene score and generic measures of constraint and generic network features further demonstrate that high-confidence ASD genes have a relationship with constraint scores in that many confirmed ASD genes are constrained against LoF variation (pLI > 0.9), and that they are likely well-studied genes (Figure 3). As genes are associated with disease, they become more studied, and they usually collect a high number of functional and physical annotations. While these annotations may be biologically relevant, they can impact GBA ML studies in a negative way by increasing the effects of multifunctionality, as discussed below.

### Overlap in the subset of genes identified as likely ASD candidate risk genes

We next examined whether overlap among top ranked genes may still exist despite low overall correlation (Table 2). For example, while forecASD and Princeton share 831 genes in their top rankings, forecASD is able to recover 118/144 SFARI high-confidence genes from a potential 1 803 compared to the 83/144 recovered from a potential 2 467 by Princeton (Table 4, red highlights). Likewise, Princeton and ExAC pLI share 1 045 genes in their top rankings, but ExAC pLI captures 121/144 from a potential 3 220 (Table 4, red highlights). This again shows that the systems-based ML studies are not performing as well as those with ASD-specific genetics information, and that they are providing little ASD-specificity above that provided by the generic measures of constraint.

We noted that multiple genes identified in previous TADA analyses are no longer statistically significantly associated with ASD in Satterstrom (i.e. only 36 of iHart’s significant findings are in Satterstrom’s 102) (Table 4 and Supplementary Table 1), and that the TADA analyses only share 17 genes in their top findings (Supplementary Table 2). There are seven genes found by recent TADA studies, which at the time of their publications, were considered novel findings; however, they are now considered to be SFARI-HC genes (Supplementary table 3). The differences in overlap of top findings between the TADA analyses further highlights that the differences between the underlying models need to be investigated more closely.

### Feature importance in forecASD algorithm

While forecASD had significantly better performance with SFARI and novel high-confidence ASD risk genes compared to other systems-based GBA ML studies, it is performing with low precision (P20R_forecASD_=4.63-11.21%) (Table 2, Table 3). If a GBA ML method is to be considered successful, it must be able to generalize to new data, and highly rank true positives. To understand the driving force behind forecASD performance, we examined its performance using training feature sets made up of different combinations of the original features used in the model (Table 5).

**Table 5:**
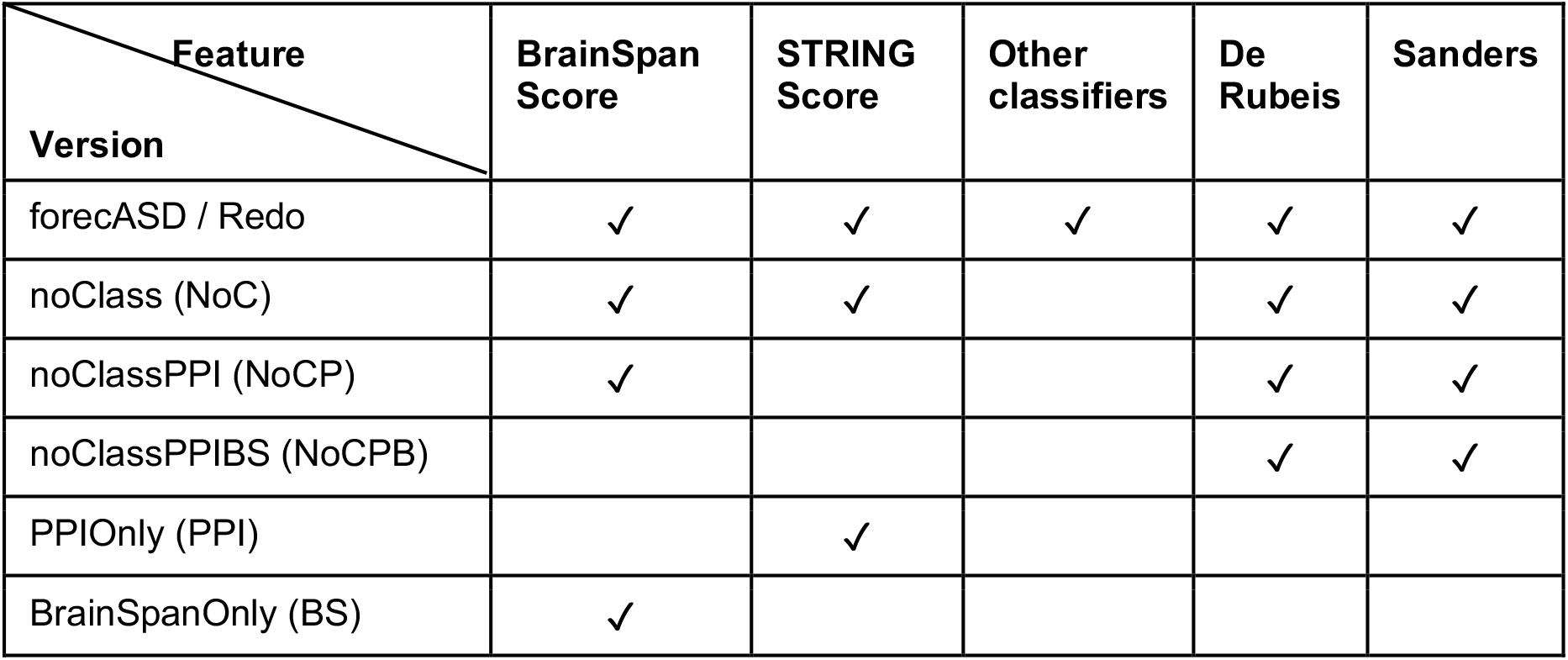
Features included in the different forecASD analyses.

| **Feature**<br>**Version** | **BrainSpan**<br>**Score** | **STRING**<br>**Score** | **Other**<br>**classifiers** | **De**<br>**Rubeis** | **Sanders** |
| --- | --- | --- | --- | --- | --- |
| forecASD / Redo | ✓ | ✓ | ✓ | ✓ | ✓ |
| noClass (NoC) | ✓ | ✓ |  | ✓ | ✓ |
| noClassPPI (NoCP) | ✓ |  |  | ✓ | ✓ |
| noClassPPIBS (NoCPB) |  |  |  | ✓ | ✓ |
| PPIOnly (PPI) |  | ✓ |  |  |  |
| BrainSpanOnly (BS) | ✓ |  |  |  |  |

Other classifiers included as features in this analysis were DAWN, Princeton and DAMAGES. The features from the DAWN algorithm were their list of risk ASD genes (rASD), and network score and minimum FDR. The features from the DAMAGES algorithm were the D and Ensemble scores. There were four FDR values from Sanders: tada_asc+ssc+del (most thorough with exome data and small *de novo* dels), tada_asc+ssc (both sources of exome data), tada_asc (ASC exome only), and tada_ssc (SSC exome only). The per-gene Bayes Factor from DeRubeis was also included (TADA_BF).

We evaluated the forecASD models on both novel-HC and SFARI-HC gene sets. In both evaluations, we found that forecASD versions incorporating genetics information had significantly better performance than the versions using only protein-protein interaction or BrainSpan gene-expression data (Figure 5, Supplementary tables 4 and 5). Notably, we found that the forecASD version incorporating only genetics data (noClassPPIBS) had overlapping 95% confidence intervals for precision at 20% recall of novel and SFARI-HC genes with the full forecASD version (i.e. P20R_novelHC:forecASD_=4.63-11.21%; P20R_novelHC:noClassPPIBS_=2.88-9.73%) (Figure 5, Supplementary tables 4 and 5). These results show that forecASD performance is driven by genetic association data (Figure 4; we note that in the forecASD preprint, STRING was considered the most informative feature, but we were unable to reproduce this result with their code despite reproducing their classification results; we believe it is an error). Taken with our other findings, the implication is that supplementing ASD-specific genetics data with heterogeneous biological data is likely not useful for disease gene discovery, especially when considering the unknown reliability and biases within the data.

**Figure 4:**
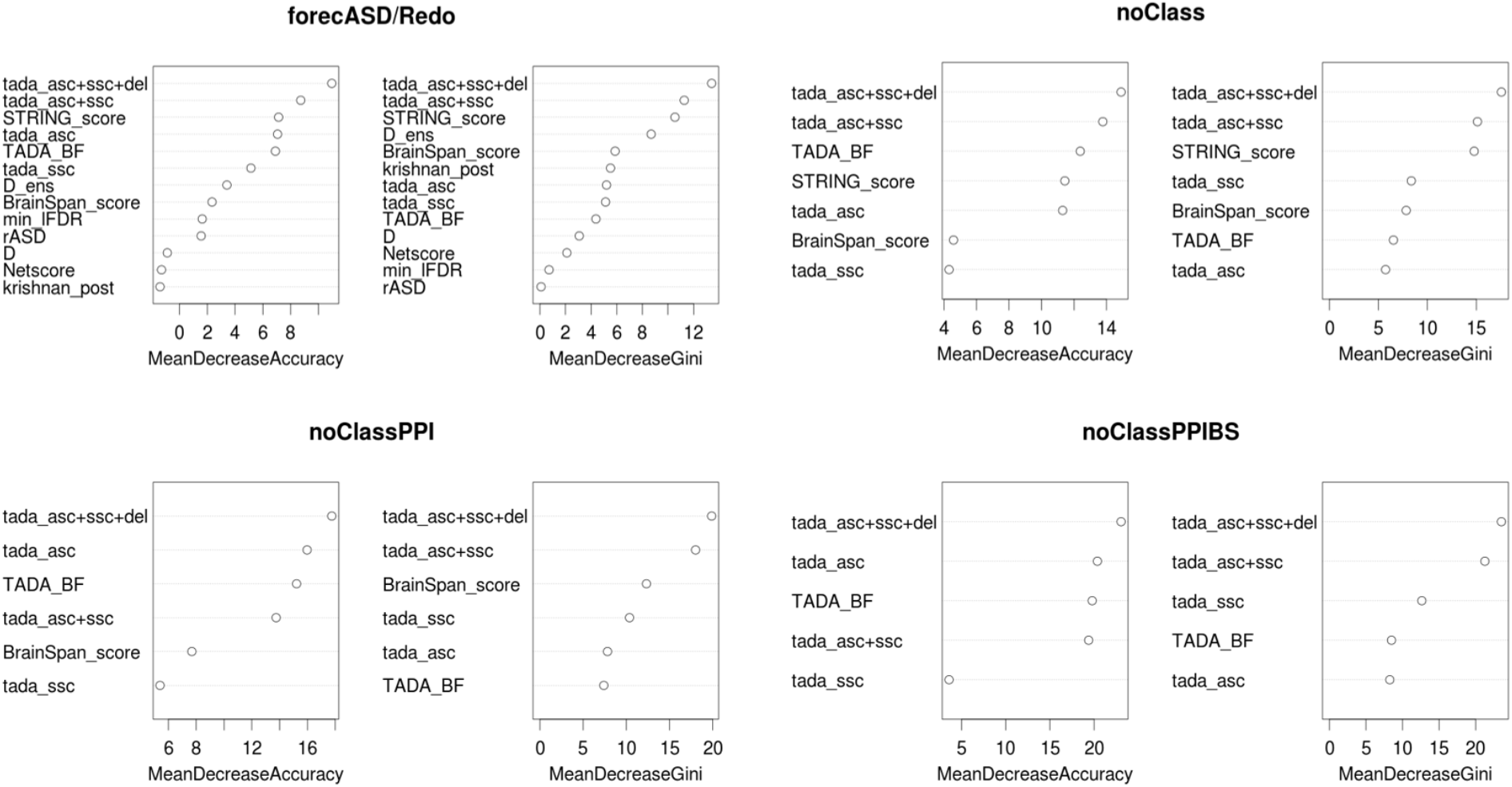
Genetics features drive forecASD performance. The most comprehensive score from the Sanders, tada_asc+ssc+del, was ranked as the most important feature for discerning ASD from non ASD training genes in each version, followed by other TADA-based statistics.

**Figure 5:**
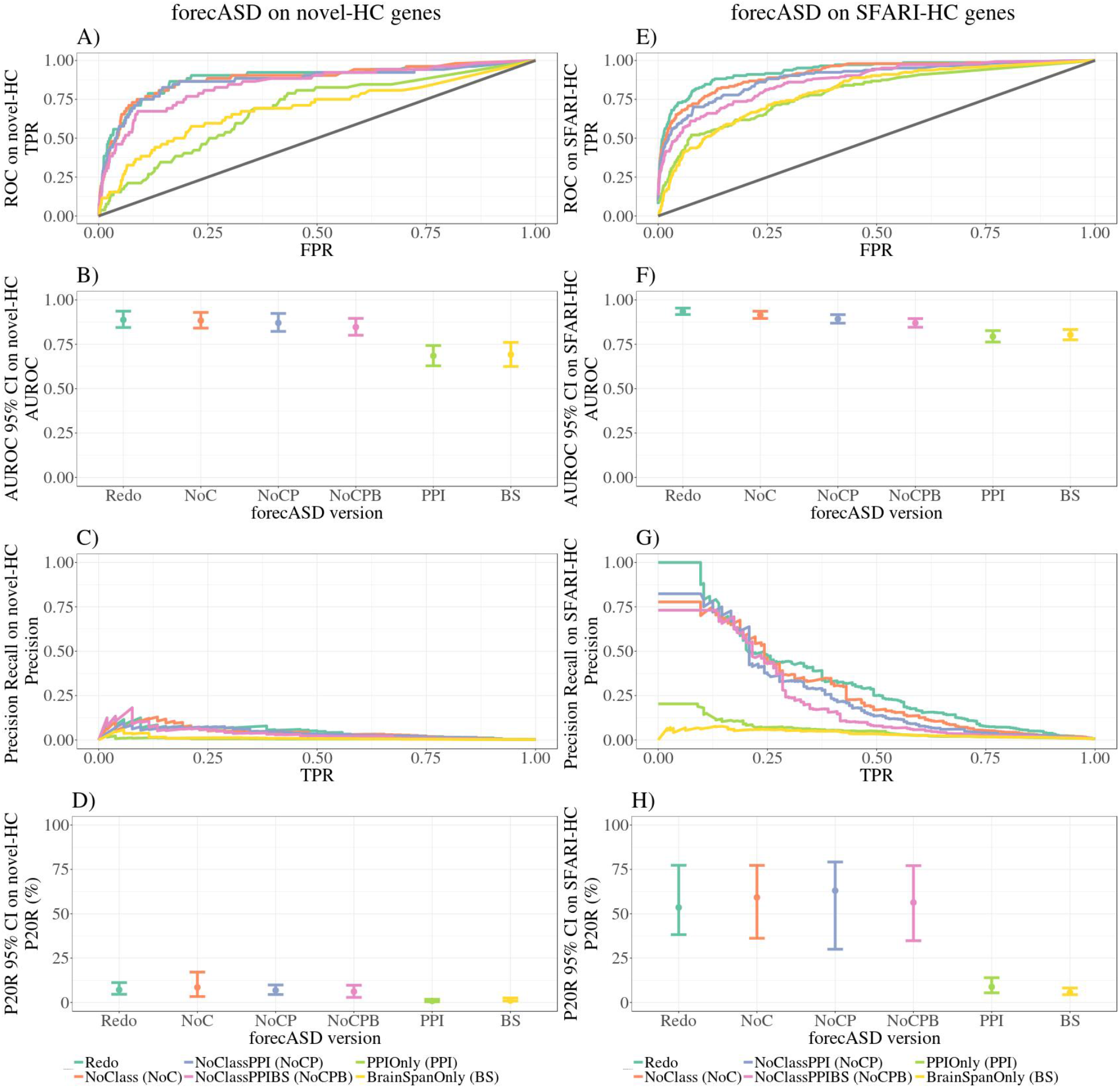
ROC, Precision-Recall and summary statistics for forecASD versions on novel-HC and SFARI-HC genes. Versions without genetics information, PPIOnly and BrainSpanOnly, show significantly worse performance in both tests. 95% confidence intervals were created from 2500 stratified bootstrap samples. TPR, True Positive Rate; FPR, False Positive Rate; AUROC, Area Under the Receiver Operator Curve; P20R, Precision at 20% Recall

## Discussion

Our investigation has shown that GBA ML methods that do not use ASD genetics information have limited utility. This appears to be because non-genetic association data provides little to no useful information above that provided by generic measures of disease gene likelihood. This finding likely has implications for other attempts to prioritize genes for complex human genetic diseases: using heterogeneous biological network data likely has diminishing returns due to poor real-world performance and biases.

### Non-equivalence of genetic association studies

A complication of our study was that the ASD genetic association studies agree poorly, even when analyzing heavily overlapping sets of subjects. For example, the recent work of Satterstrom et al., fails to replicate many of the genes considered significant ASD risk genes reported by De Rubies et al., despite using essentially all the data from De Rubies et al. The reason for this is not clear. One possibility is that many of the genes reported by De Rubies et al. were false positives uncovered by Satterstrom having more data. Arguing against this, all of the genes identified by De Rubies et al. were considered high-confidence ASD genes by SFARI Gene at the time of our analysis. Another likely culprit is that the methods for detecting statistical association of very rare *de novo* variants with phenotypes were changed substantially in Satterstrom et al. (2020). We note that iHart uses the same TADA model as Sanders, but with an increased number of samples from multiplex families, and Satterstrom and iHart show the highest agreement in ranking (R_S_=0.92) and overlap of significant genes (52/80). For this reason we consider it likely that the incorporation of pLI and MPC in Satterstrom et al. has a larger impact on the results than changes to the underlying data. Regardless, it is a caveat of our study that there is apparently no universally trustable gold standard set of ASD genes. The impact of this on the interpretation of our study is limited, because as we show, the set of genes used for evaluation does not change the performance outcomes substantially.

### ML methods are comparable to generic measures of LoF constraint

Proposed use cases of the GBA ML studies include prediction and/or prioritization of ASD risk genes, framing WES/WGS results for further exploration in resequencing or mechanistic studies, and/or uncovering new and delineating possible pathways implicated in ASD etiology (Brueggeman et al., 2020; Duda et al., 2018; Krishnan et al., 2016; Lin et al., 2018; Liu et al., 2014; C. Zhang & Shen, 2017). Overall, for GBA ML study to be considered successful in identifying novel ASD genes, it should highly prioritize known ASD genes and provide additional, specific and unbiased predictions above that which could be obtained from generic measures of constraint. We have shown that the systems-based ML studies failed to do so.

As discussed previously, we expect that methods employing GBA would tend to rank generically “disease-related” genes highly because they are well studied, highly annotated and highly connected within networks. Thus, methods biased towards generic rankings, such as PANDA and DAMAGES, likely struggle to identify novel and disease-specific relationships (Figure 3). Conversely, GBA methods which are not biased towards generic rankings, such as Princeton and FRN, may perform badly because the main source of apparent performance of GBA methods is their ability to prioritize well studied genes (“multifunctionality bias” as per Gillis and Pavlidis)(Figure 3). While we found that, overall, the system-based GBA studies perform with low precision, we also found that two studies, Princeton and FRN, are not biased towards well studied, highly annotated, highly connected genes (Table 3, Table 4, Figure 3). These two studies built complex functional interaction networks from multiple data types, including protein-protein interaction and gene expression data. They used Gene Ontology annotations to define “gold standards” of functional relationships and Bayesian frameworks for weighting and data integration (Duda et al., 2018; The Gene Ontology Consortium, 2019; Greene et al., 2015; Krishnan et al., 2016). Their poor performance could be due to their GO functional categorization not aligning well with the multiple biological data types and/or not providing useful ASD-specific information. However, it is much more likely that these studies do not perform well because due to the effects, or lack thereof, of multifunctionality bias (Gillis & Pavlidis, 2011; Pavlidis & Gillis, 2012).

While further investigation into each study is required to delineate how multifunctionality bias is affecting their performance, a consistent finding across the GBA ML studies was high agreement between the studies and generic measures of constraint against LoF variation (Figure 3). We have confirmed that measures of constraint against LoF variation are able to identify ASD genes, albeit with low precision, and that they agree with generic network features and annotations (Table 3, Table 4, Figure 3). Many previous studies have found ASD genes, particularly those with high numbers of recurrent *de novo* variants, to be enriched for genes under high evolutionary constraint, and LoF constraint has previously been reported to be positively correlated with the number of physical interaction partners (De Rubeis et al., 2014; Karczewski et al., 2019; Lek et al., 2016; Ruzzo et al., 2019; Satterstrom et al., 2020). From this, we can confirm that measures of constraint against LoF variation measure generic susceptibility to disease, and that high constraint does not automatically guarantee a particular disease status, necessitating incorporation with data specific to the disease at hand to increase precision (Cummings et al., 2019; Karczewski et al., 2019; Lek et al., 2016).

Furthermore, while these measures are also correlated with numbers of interaction partners, functions and publications, they may point towards more biologically relevant information, such as the ability of a gene to influence different phenotypic traits, rather than number of connection partners based on network structure (“hubness”) (Gillis & Pavlidis, 2011; Pavlidis & Gillis, 2012).

The implication of this analysis is that supplementing ASD-specific genetics information with measures of constraint may provide a more fruitful avenue forward compared to creating GBA ML methods using biased biological networks. We can see this already being done by the Satterstrom TADA analysis by their incorporation of the pLI and MPC into the method in attempts to provide more detailed information about variant classes with higher burden in ASD probands (Satterstrom et al., 2020).

In summary, our results demonstrate that despite using complex data and sophisticated algorithms, ASD GBA ML methods fail to outperform generic measures of disease gene likelihood such as pLI. We suspect this is likely to generalize to the study of other genetic disorders.

## Acknowledgement

This work was supported by the SFARI Foundation and an NSERC Discovery Grant. MG was supported in part by a scholarship from the UBC Bioinformatics Graduate Program via the NSERC CREATE program in High-Dimensional Biology and a CGS-M scholarship.

## Data availability

Code and publicly available raw data used in this analysis are available on request.

## Supplementary text

Here we provide more detailed descriptions of the gene prioritization methods we examined in this study, and summarize the performance evaluations and claims from the source publications.

### Genetic association approaches

TADA: An increasingly common test for association between rare genetic variation and disease, particularly in the field of autism genetics research, is the Transmission and *De Novo* Association Analysis (TADA) test (De Rubeis et al., 2014; Feliciano et al., 2019; He et al., 2013; Ruzzo et al., 2019; Sanders et al., 2015; Satterstrom et al., 2020). TADA employs a gene-level approach by allowing for recurrence of multiple types of variants to be collapsed in order to maximize power to find risk genes (He et al., 2013). The test uses data from *de novo* and/or inherited variants identified by large scale sequencing studies of simplex and multiplex families and case-control cohorts. Using this data, TADA builds a likelihood model based on allele frequency, relative risks of different classes of variation, and mutation rates to estimate a gene’s likelihood of being involved in the phenotype. TADA can be seen as a “family of methods” (fundamentally, statistical models) because it can be applied using only *de novo* variation (TADA-Denovo) or using *de novo*, inherited and case-control variation (TADA), and requires parametrization of multiple variants (He et al., 2013). One implication of the lack of well-established and validated methods for gene-level association studies of rare variants it that two studies of the same cohort can get different results, even if they both use a method labeled “TADA.” Most recent ASD sequencing studies employing a TADA test provide an association score for each gene in the genome, and identify a subset a genes significantly associated with ASD under their model, at some expected false discovery rate (De Rubeis et al., 2014; Ruzzo et al., 2019; Sanders et al., 2015; Satterstrom et al., 2020). The genome-wide association scores allow us to compare prioritization of ASD risk gene candidates based on genetic association to other genome-wide prioritization scores based on other types of non-genetics data.

Iossifov LGD: First, they calculated a gene’s “vulnerability score” based on: 1) A likelihood model of expected LGD variants in a typical gene built from synonymous variation data in the parents of the SSC and the control neurotypicals; and 2) The proportion of causal ASD genes estimated from the ascertainment differential for LGD variation between affected probands and unaffected sibling controls (Iossifov et al., 2015). By combining the “vulnerability score” with the observed load of LGD variants in proband WES data, they created a heuristic prioritization score for ASD genes (Iossifov et al., 2015).

### Princeton

(Krishnan et al., 2016) used a support vector machine (SVM) trained on a human brain-specific functional interaction network built from multiple protein-protein interaction databases, gene expression datasets, and other regulatory and genetic and chemical perturbation data (Greene et al., 2015). Their training labels included 549 positive genes weighted by the strength of evidence of association with ASD (E1,2,3,4), and a set of 1189 manually curated non-mental health disease genes. Their feature space was a gene-gene matrix where the cells represented the probability of a gene-gene interaction calculated from their brain-specific functional interaction network. Using their gene-gene matrix, and their ASD-positive and ASD-negative training gene sets, they fit a linear SVM with a penalty parameter to control misclassification of their evidence-weighted labels (i.e. lower misclassifications of E1, high-confidence labels). They ran 5-fold cross-validation 50 times on different subsets of their evidence-weighted training labels and found that the model with all evidence-weighted labels had the best performance for separating positive and negative genes. In theory, the SVM fit a linear plane in the high-dimensional feature space which was able to maximize the separation between positive and negative training genes. For each candidate gene, the distance between the candidate gene and discriminant hyperplane (i.e. prediction from computing the linear kernel function) was converted to a probability using regression; an average probability was taken across each of the 5 cross-validation folds. Prediction scores were provided for 25 825 genes, and they identified their top decile of genes as likely ASD risk gene candidates. Their published evaluation and validation of their ranking system included: 1) Calculating enrichment of genes with *de novo* mutations in independent ASD sequencing studies in their top decile (Sanders et al. (2012), O’Roak et al. (2012), Iossifov et al. (2012), Neale et al. (2012), Iossifov et al. (2014), and De Rubeis et al. (2014)); 2) Calculating enrichment of experimentally determined targets of ASD-related proteins and pathways, such as FMRP and MAPK signalling, in their top decile; and 3) Calculating enrichment of genes found to be associated with intellectual disability, schizophrenia, and other developmental disorders in their top decile. From their main evaluation, they found that their evidence-weighted labels had significantly better performance during cross validation than other combinations of training labels, and that there was significant enrichment of genes found to have *de novo* likely damaging variation in independent ASD sequencing studies in their top decile of genes. Overall, they concluded that their method was able to prioritize many new ASD candidate risk genes, and claimed that their top ranked genes had the potential to speed up ASD gene discovery.

### FRN

(Duda et al., 2018) used a random forest classifier with a brain-specific functional interaction network. They built their network from human, rat and mouse gene expression datasets from non-cancer related brain experiments, multiple protein-protein interaction databases, and protein docking and phenotype annotations. They utilized 143 ASD genes from SFARI 1, 1S, 2, 2S and Sanders as positive training labels, and 1176/1189 of the negative non-mental health genes from Princeton. Their feature space was a gene-gene matrix where the cells represented the probability of a gene-gene interaction based on their brain-specific functional interaction network. Using their gene-gene matrix, and their ASD-positive and ASD-negative training gene sets, they trained 5 different machine learning models with 5-fold cross-validation, and found that their random forest model had the best performance based on the average AUROC from the 5-folds. Random forests are built from multiple decision trees which segment the feature matrix into a number of simple regions by recursive binary splitting. In each decision tree of a random forest, each split in each tree uses a random sample of features, and at each successive split, the best splitting rule is chosen so that the two new regions are as pure as possible. In other words, the feature and its threshold which give the best separation between the positive and negative training data is chosen at each split point in each tree. The leaves at the bottom of a decision tree are called terminal nodes. Predictions are made for candidate items (genes) based on which decision path it follows, and the proportion of positive and negative training observations in the terminal node. In other words, after allowing a candidate gene to follow a decision path and entre a terminal node, if the majority of the genes in the terminal node are positive training genes, the candidate gene will be predicated as a positive. Prediction scores were provided for 21,114 genes, and they identified their top 2,111 genes as likely ASD risk gene candidates. Their published evaluation and validation of their ranking system included: 1) Calculating the enrichment of genes with recurrent and non-recurrent *de novo* LoF mutations in ASD probands and unaffected siblings from the SSC (Iossifov et al. (2014)) and MSSNG (Yuen et al. (2017)) in their top decile; and 2) Calculating the enrichment of genes found to be involved in Alzheimer’s disease, Parkinson’s disease and ataxia. From their evaluation, they found significant enrichment of genes with *de novo* LoF mutations in SSC and MSSNG probands in their top decile, and an absence of significant enrichment for genes involved in other brain-related disorders. Overall, they concluded that their method predicted genes with evidence of ASD association and was able to propose numerous novel genes they claimed had a high likelihood of contributing to ASD.

### DAMAGES

(C. Zhang & Shen, 2017) used cell-type specific expression data from 24 mouse central nervous system cell types from 6 regions, and measures of constraint against LoF and missense variation from ExAC to try to identify ASD risk genes. First, they created a DAMAGES (D) score built from gene expression profiles of 145 genes found to have *de novo* LGD variants in probands and unaffected siblings from Iossifov et al. (2012), Neale et al. (2012), O’Roak et al., (2012) and Sanders et al. (2012) using Principal Component Analysis (PCA). Regression analysis was used to evaluate how each principal component from the PCA analysis was able to predict a gene’s variation source as proband or sibling control. Next, they used logistic regression to combine the D score with measures of constraint against LoF and missense variation to create an ensemble score (E). Their training labels for their logistic regression classifier were 36 genes found to have 2 or more *de novo* LGD mutations in ASD probands, and 156 genes with only 1 or more *de novo* LGD mutations in sibling controls. Their feature space consisted of the D score (PCA-based gene expression profiles) and ExAC constraint scores. They used logistic regression to estimate the effect size of each feature, and then calculated an ensemble (E) score for each candidate gene predicting its likelihood of being a haploinsufficient ASD gene. The mouse genes they used to create the D score were mapped to human orthologs so the constraint scores could be added to create the E score. They identified the top 117 genes by E score as likely ASD risk gene candidates. We kept E scores for 15 881 genes with single, unambiguous mappings to human genes. Their published evaluation and validation of their ranking system included: 1) Calculating enrichment of genes with LGD mutations from sequencing studies published after 2012 (De Rubeis et al. (2014), Iossifov et al. (2014)), and 438 SFARI genes by category (S,2,3,4,5,6) in the top ranking of the D score; 2) Comparing the D score and Ensemble score to constraint measures alone, and ranks provided by Princeton by calculating a modified precision recall statistic. From their evaluations, they found enrichment of genes found to have *de novo* likely damaging variation in independent ASD sequencing studies, and that their method have favourable performance compared to other studies. Overall, they concluded that their gene expression signatures reflected haploinsufficiency in ASD, and claimed that it was able to predict whether or not likely damaging variants confer increased risk to ASD.

### RF_Lin

(Lin et al., 2018) employed a random forest classifier using gene-level constraint measures from ExAC and a weighted network built from BrainSpan and InWeb protein-protein interaction data as features (Li et al., 2017; Miller et al., 2014). Their training labels are the same employed in the FRN method above. See ASD_frn for description of random forests. In theory, their random forest was able to split the feature space (constraint, weighted network information) into regions which could separate their positive training genes from the negative training genes, and thereby predict which candidate genes were the most similar to positive training genes. Prediction scores were provided for 17 099 genes, and they identified their top 2 089 genes as likely ASD risk gene candidates. They did not provide scores for their training labels. Their published evaluation and validation of their ranking system included: 1) Calculating enrichment of genes found to have *de novo* LoF or missense mutations from 2517 SSC families (Iossifov et al. (2014)) and MSSNG (Yuen et al. (2017)) in their top decile; 2) Comparing their ranking system to ExAC pLI, Iossifov, Sanders, Princeton and DAMAGES by calculating decile enrichment of 130 SFARI category 3 genes, and 43 genes with recurrent *de novo* LoF mutations identified in Stessman et al. (2017), Wang et al., (2016), Yuen et al., (2017), and Li et al., (2017); and 3) Comparing their ranking system to ExAC pLI, Iossifov, Sanders, Princeton and DAMAGES by calculating the AUROC with their labelled and unlabelled data, with the 130 SFARI category 3 genes, and with the 43 genes with recurrent *de novo* LoF mutations. From their evaluations, they found significant enrichment of genes with *de novo* LoFs in SSC and MSSNG probands, and that their method showed higher enrichment of 173 candidate ASD genes in their top decile compared to other methods. Overall, they concluded that their method demonstrated that spatiotemporal gene expression and constraint metrics predicted ASD risk genes, and claimed that their method provided many new candidate genes with strong evidence of contributing to ASD.

### PANDA

(Y. Zhang et al., 2020) is a graph neural network type classifier. They built an unweighted and undirected human molecular interaction network from experimentally documented physical protein interactions using data from a previously established protein-protein interaction network (Menche et al., 2015), and BioGRID (Oughtred et al., 2019). Their training labels included 760 autism-associated genes from SFARI Gene 2.0 (Abrahams et al., 2013), and Online Mendelian Inheritance in Man (Hamosh et al., 2005) weighted by confidence of association (0.5, 0.75, 1.0), and 1102 non-autism associated genes from FRN and Princeton above. They training their algorithm using five-fold cross-validation. The human molecular interaction network was represented as a graph whereby the genes were nodes, and their interactions were edges. They described the local structural properties of each node using six node centrality measures and calculated how often a node appeared in 69 different orbits (distinct positions of nodes) in 4- and 5-node graphlets (connected, nonisomorphic induced subgraphs). Next, a local transition function aggregated all the network properties of each node, and its direct neighbours in ten dimensions to obtain node embeddings (10-dimenstional vector describing each node). Using these 10-dimensional spatial representations, a global output function was used to predict the class label of each node (1=autism-associated, 0=no autism-association). In addition, they used a sigmoid function to transform each prediction into a probability and computed a loss function to penalize embeddings that encode neighbours very differently. They had four layers to their classifier, meaning their local transition and global output functions were implemented as four feed-forward neural networks. Prediction scores were provided for 23 472 genes, and they identified an “autism subnetwork” of 2 346 genes. Their published evaluation and validation of ranking system included: 1) Evaluation of the classification performance of PANDA by calculating sensitivity, specificity, classification accuracy, precision@k (proportion of positive autism-genes in top-k ranked list), and comparing to three other types of machine learning algorithms; 2) Calculating enrichment of genes with *de novo* likely disrupting mutations from probands and their unaffected siblings an independent sequencing study (Iossifov et al., 2014) in their top decile of genes; 3) Calculating the specificity of PANDAs rankings to ASD by looking at the distribution of Alzheimer’s, Parkinson’s and Epilepsy genes in their rankings; 4) Investigating their unlabelled top-ranked genes for potential association with ASD; and 5) Identifying an “autism subnetwork” from the human molecular interaction network made up of 2 346 genes. From their evaluations, they found that PANDA was the best performing type of machine learning algorithm, and that their top decile of genes from ranked gene list showed significant enrichment for recurrent and non-recurrent proband DN-LGDs. Further, they found that their rankings specific to ASD due to lack of significant enrichment of other disease genes in their top decile. Overall, they concluded that PANDA was able to search the graph space to find genes with similar topological properties or indirect connections to known autism genes, and predict which are the most likely to be autism genes themselves.

### Hybrid Genetics-GBA machine learning studies

The studies in this section used a combination of ASD-specific features and other features to build their models. The ASD-specific features come from genetic association data in the studies described above. Information from the two classes of features are integrated prior to training a machine learning algorithm to distinguish ASD from non-ASD risk genes, using high-confidence ASD genes from genetic association studies as their positive training set.

### DAWN

(Liu et al., 2014) targeted their search to genes found to be expressed in the prefrontal and motor-somatosensory (PFC-MSC) neocortex during the 10-24 weeks post-conception phase based on previous findings from Willsey et al.(2013). Willsey et al., (2013) found that the PFC-MSC from 10-24 weeks post-conception was a potential nexus for risk based on coalescence of gene expression during that time. They built a co-expression network from BrainSpan data of the selected regions and time points using Weighted Gene Co-expression Network Analysis (WGCNA) and overlaid genetic association statistics from a TADA model to identify ASD risk gene candidates (Langfelder & Horvath, 2008; Liu et al., 2014). The TADA scores utilized were calculated from rare *de novo* likely damaging variants, rare transmitted likely damaging variants and case-control likely damaging variants from multiple sources, including Iossifov et al. (2012), Neale et al. (2012), O’Roak et al. (2012), Sanders et al. (2012), and the ARRA Autism Sequencing Consortium, among others. They used unsupervised model-based clustering and a hidden Markov random field to model the correlation of genetic association scores across the co-expression network to identify co-expressed nodes with high genetic evidence of association with ASD (“network ASD genes”). Next, they used a false discovery rate procedure to determine which of the “network ASD genes” were most likely to contribute to ASD (“risk ASD genes”, rASD genes). They provided prediction scores for the 10 233 genes in the network which had exome data. They identified their top 127 genes (FDR < 0.05) to be likely ASD risk gene candidates. Their published evaluation and validation of their ranking system included: 1) Two permutation dilution experiments were conducted whereby the signal from genetic association or co-expression data was diluted to determine the sensitivity of the rASD gene list to either signal; and 2) Calculation of enrichment of *de novo* LoF mutations identified in a targeted sequencing study of 44 ASD candidate genes in 2448 ASD trios in their 127 rASD genes. From their evaluation experiments, they found that DAWN was sensitive to both the TADA signal and the co-expression signal, and that DAWN was able to identify genes found to have more *de novo* LoF variants in ASD probands. Overall, they concluded that there was a high likelihood that DAWN had identified true ASD risk genes.

### forecASD

(Brueggeman et al., 2020) is a stacked random forest ensemble classifier. They built the first layer from BrainSpan spatiotemporal gene expression data (Miller et al., 2014) and a protein-protein interaction matrix built from STRING (Szklarczyk et al., 2019). The second layer was built from the scores from layer 1, and scores from other studies, all of which are included in my study (Princeton, DAWN, DAMAGES, De Rubeis and Sanders). Their training data included 76 SFARI high-confidence ASD genes, and 1000 random non-SFARI genes. See ASD_frn for description of random forests. In theory, each random forest layer they built was able to split the feature space (BrainSpan, STRING, genome-wide ASD prediction sores) into regions which could separate their positive from their negative training genes, allowing for candidate gene predictions to be made based on shared/similar associations to the positive training genes in the feature space. Prediction scores were provided for 17 957 genes, and they identified their top 1787 as likely ASD risk gene candidates. Their published evaluation and validation of their ranking system included: 1) Calculating enrichment of genes with *de novo* likely disrupting mutations in MSSNG (Yuen et al., (2017)) and Spark (unpublished at time of forecASD development) probands, SFARI 3, 4, 5 and syndromic genes, and gene targets of CHD8 and FMRP in their top decile of ranks; 2) Calculating the AUROC for their score, and scores from Princeton, DAMAGES and Sanders on SFARI category 1 and 2 genes, and SFARI category 3 genes; 3) Calculating enrichment of genes with *de novo* likely disrupting mutations in MSSNG (Yuen et al. (2017)) and Spark (unpublished at time of forecASD development) probands in the top decile of the scores from Princeton, DAMAGES and Sanders for comparison; and 4) Fitting logistic regression models to assess how much forecASD is adding to genetic TADA signals. From their evaluations, they found their method was better able to classify SFARI 1 and 2 genes and SFARI 3 genes, and they showed greater enrichment in their top decile for genes found to have recurrent *de novo* likely damaging variants in Spark and MSSNG probands compared to other studies. Further, they concluded that forecASD was able to provide biological context to TADA genetic signals important for prioritization. Overall, they concluded that their method was able to generalize to new data and claimed that they had created a valuable framework for combing both genetics and non-genetics data to prioritize ASD risk gene candidates which will be useful for when ASD gene discovery by genetic association slows.

